# caRBP-Pred: Deep Learning-based Prediction of Chromatin-Associated RNA-Binding Proteins Using Short Peptide Sequences

**DOI:** 10.64898/2026.01.12.698323

**Authors:** Qiang Sun, Feng Yang, Hao Sun, Xiaona Chen, Huating Wang

**Affiliations:** Department of Orthopaedics and Traumatology, Li Ka Shing Institute of Health Sciences, The Chinese University of Hong Kong, Hong Kong, China; Center for Neuromusculoskeletal Restorative Medicine, Hong Kong Science Park, Hong Kong SAR, China; Warshel Institute for Computational Biology, Faculty of Medicine, Chinese University of Hong Kong, Shenzhen, Guangdong, China

**Keywords:** caRBP-Pred, chromatin-associated RNA-binding proteins, deep learning

## Abstract

RNA-binding proteins (RBPs) are pivotal in cellular processes ranging from RNA metabolism to 3D genome organization. A distinct subset, chromatin-associated RBPs (caRBPs), binds directly to chromatin to function as transcriptional regulators. However, identifying caRBPs via traditional methods like Chromatin Immunoprecipitation Sequencing (ChIP-seq) and Mass Spectrometry (MS) is labor-intensive and costly. While computational tools for DNA- and RNA-binding protein (DRBP) prediction exist, they often rely on outdated Gene Ontology annotations and fail to capture the unique characteristics of chromatin association. Here, we introduce caRBP-Pred, a novel deep learning approach combining Convolutional Neural Networks (CNN) and Bidirectional Long Short-Term Memory networks (BiLSTM). Unlike previous methods utilizing full-length sequences, our model is trained on chromatin-contact peptides derived from mouse embryonic stem cells (mESCs). caRBP-Pred achieves a superior Area Under the Curve (AUC) of 0.81 using peptide sequence information alone, significantly outperforming existing DRBP predictors which exhibit low recall for caRBPs. We further predicted 52 potential caRBPs in mice. Notably, validation against human homologs confirmed that our model accurately predicts candidates with experimentally verified chromatin-binding capabilities. Collectively, caRBP-Pred is the first tool specifically designed to predict caRBPs based on chromatin-contact peptides, offering a valuable resource for investigating regulatory roles of caRBPs on transcription.

## 1 Introduction

RBPs are well-established as key regulators of gene expression, operating through diverse mechanisms. Their classical role in post-transcriptional regulation encompasses RNA alternative splicing, localization, translation, and stabilization (1–5). More recently, RBPs have also been implicated in transcriptional regulation and 3D genome organization (6–11). Thus, RBPs can profoundly influence a wide range of biological functions through multiple pathways. Consequently, a comprehensive analysis of RNA/DNA-RBP complexes and their molecular functions is essential for a deep understanding of these interactions. Recent studies have revealed that a wide variety of RBPs can bind to open chromatin and active chromatin regions including promoters and enhancers. These interactions influence transcription and/or mediate co-transcriptional processing, providing novel insights into the field of gene expression (6,7,12,13). For example, QKI5, which is a caRBP that identified in K562, 293T, and THP1 cells, can bind to chromatin independent of RNA or transcription factors, and can function as a transcriptional activator to activate the monocytic differentiation-associated genes(12). Another caRBP, Dazl, co-localized with PRC2 at the promoters of genes in mESCs and is involved in the formation of phase-separated PRC condensates(14). Currently, numerous experimental methods have been used to identify the DNA binding sites of RBPs, such as ChIP-seq (7), CUT&RUN, CUT&Tag, and mass spectrometry (2,7,11,12,15). However, the identification of caRBPs using these methods is very time-consuming and costly, due to the complex experimental procedures involved. Moreover, they are limited in their capacity to identify multiple RBPs simultaneously. Over the past decades, a suite of computational methods leveraging deep learning and machine learning has emerged to predict RBPs. These models are typically trained on protein or RNA primary sequences, often incorporating derived features such as physicochemical properties (e.g., hydrophobicity, polarity, and side chain charge) and secondary structures. For example, tools like RBPPred and NAbind utilize physicochemical properties or electrostatic information with Support Vector Machines (SVM) to predict RBPs(16,17). Other models, such as Deep-RBPPred, harness Convolutional Neural Networks (CNNs) to enhance prediction accuracy(18). Additionally, models like rBPDL integrate CNN-LSTM architectures with full-length protein inputs to further improve performance(19).

However, a critical distinction exists between generic DRBPs and caRBPs. While DRBPs are typically defined by their capacity to bind directly to specific DNA or RNA sequences, caRBPs represent a distinct functional subset. caRBPs associate with chromatin not only through direct nucleic acid binding but also potentially via interactions with histone modifications, chromatin-associated proteins, or through phase separation mechanisms(14). Unlike classical DRBPs, caRBPs are characterized by their ability to bind chromatin in an RNA-independent manner to regulate transcription and 3D genome organization. Consequently, existing DRBP predictors (e.g., iDRBP_MMC, DeepDRBP-2L, DeepMC-iNABP) face limitations: they rely on GO annotations that may lag behind current knowledge, and they are not designed to capture the unique features of chromatin-caRBP interactions(20–22).

In this study, we developed a CNN-BiLSTM model to predict caRBPs in mice, capable of capturing contextual sequence relationships and extracting deeper features than traditional models. Leveraging an existing dataset of caRBPs from mESCs that bind chromatin in an RNA-independent manner, our model integrates full-length protein sequences and chromatin-contact peptide sequences as inputs. We found that the CNN-BiLSTM model trained on peptide sequences alone displayed the best performance, whereas existing DRBP predictors showed a limited recall rate for caRBPs. Based on these results, we predicted 52 potential caRBPs in mice cells. This is the first computational tool specifically designed for predicting caRBPs using chromatin-contact peptides, providing a reliable candidate list for future investigation of RBP’s regulatory role on transcription.

## 2 Material and method

### 2.1 Data preprocessing and input representation

We assembled a comprehensive dataset comprising 403 caRBPs and 1694 non-caRBPs from mESCs, as previously reported (14). For raw protein or peptide sequences, each amino acid in a sequence was encoded into a 26-dimensional one-hot vector based on a custom mapping that accounting for standard and modified residues. Sequences were padded or truncated to a fixed length of 1000 residues using post-padding to ensure uniform input dimensions. The dataset was split into training (80%) and testing (20%) sets using a random state of 101 for reproducibility.

### 2.2 Independent dataset

For the mouse RBP dataset, we sourced a total of 2897 RBPs from the RBP2GO database (23). To ensure independence, we excluded any RBPs already present in our training and testing datasets, resulting in a final independent dataset of 1559 RBPs for further prediction analysis.

### 2.3 Prediction of protein and peptide secondary structure

To predict the secondary structures of proteins and peptides, we employed PS4 with default parameters (24). This tool generated secondary structure annotations corresponding to the primary amino acid sequences of each protein and peptide in our datasets.

### 2.4 CNN-BiLSTM model

The model combines convolutional and recurrent layers to capture both local spatial patterns and long-term temporal dependencies in protein sequences, as illustrated in Figure 1. The detailed architecture is as follows: the first component consists of two consecutive 1D convolutional layers with 32 filters each, a kernel size of 3, and ReLU activation functions. Each convolutional layer is followed by a max-pooling layer with a pool size of 2 and a dropout layer with a rate of 0.5 to prevent overfitting. The CNN module serves to extract local spatial features from the protein sequences. The output from the second convolutional block is passed through a TimeDistributed(Flatten()) layer to prepare features for sequential processing. This is followed by a bidirectional LSTM layer with 70 units per direction (140 total), configured to return full sequences. The sequence outputs are then processed by a second bidirectional LSTM layer with 70 units per direction, which returns only the final hidden state. The BiLSTM layers are designed to capture long-range bidirectional dependencies within the sequence. The final component is a fully connected output layer with a single neuron and a sigmoid activation function for binary classification

**Figure 1.**
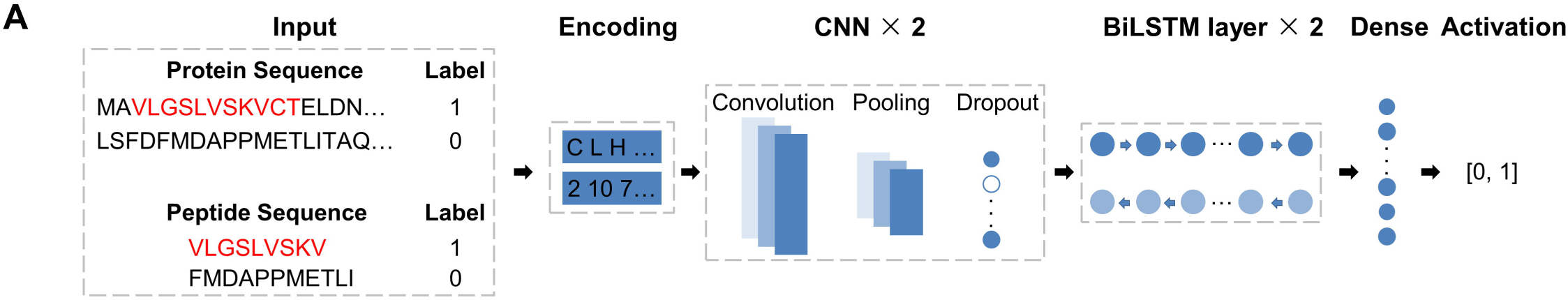
Model architecture of CNN-BiLSTM for caRBP prediction. **(A)** Flowchart of the CNN-BiLSTM model.

The primary sequences are first encoded using a one-hot encoding layer. The CNN layer, comprising convolutional, max-pooling, and dropout operations, captures the features of both sequence and secondary structure information. Subsequently, the Bi-LSTM layer extracts contextual information from these features. A dense layer then combines outputs from various convolutional kernels, followed by an activation layer that computes a score between 0 and 1 for each input sequence, indicating the likelihood of being a caRBP.

### 2.5 Model Training Parameters

The model was compiled with the Adam optimizer using binary cross-entropy as the loss function. Training was conducted for a maximum of 60 epochs with a batch size of 32. To address class imbalance, class weights of {0: 1, 1: 10} were applied, where 0 and 1 represent positive and negative peptides, respectively. An early stopping callback was implemented to monitor validation loss with a patience of 10 epochs, restoring the best weights when triggered.

### 2.6 Support Vector Machine (SVM), Random Forest (RF), and eXtreme Gradient Boost (XGBoost) model

To compare the performance of CNN-BiLSTM with other learning models, we selected the SVM, RF, and XGBoost, and the parameters for these models were set to default during the training process.

### 2.7 Motif enrichment analysis

To identify enriched motifs in the positive peptide sequences, we utilized the XSTREME module from the MEME suite (25), with negative peptide sequences serving as the background. We specifically focused on protein motifs from the Eukaryotic Linear Motif (ELM) database (version 2024) to provide biological context to our findings(26).

### 2.8 Performance evaluation

We employed a 5-fold cross-validation procedure to rigorously evaluate our deep learning model. Performance metrics were calculated by averaging the model’s performance across the five validation sets. This approach ensured that our model’s predictive accuracy was robust and generalizable across different subsets of the data. The following metrics were used for quantitative assessment of the model’s performance:

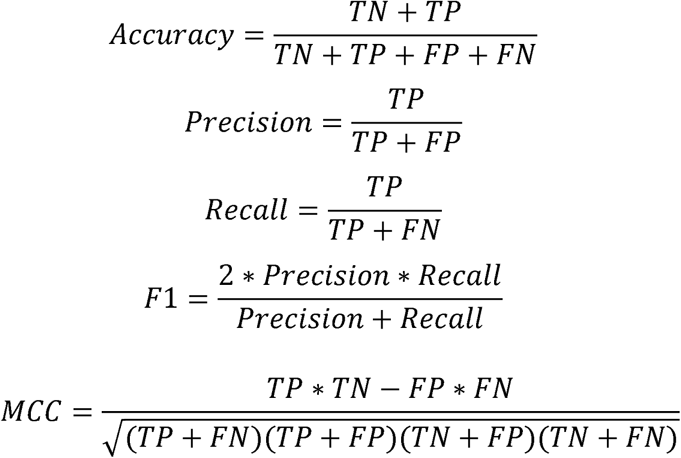

### 2.9 DRBP predictors evaluation

Since the source codes for previous DRBP predictors (iDRBP_MMC, DeepMC-iNABP, and DeepDRBP-2L) are not publicly available(20–22), we evaluated their accuracy on the caRBP training dataset using their provided web servers or executables.

### 2.10 caRBP-Pred implementation

To predict potential caRBPs, we developed an in-house script named caRBP-Pred. The script employs a sliding window approach with a window size of 50 amino acids (AA) and a stride of 1 AA to identify positive peptides within an input RBP sequence. The window size was specifically set to 50 AA to correspond with the maximum length of chromatin-contact peptides identified in our dataset, ensuring the model captures the complete sequence features of potential chromatin-binding domains. A stride of 1 AA was utilized to achieve high-resolution scanning and prevent missing motifs located at window boundaries.

## 3 Results

### 3.1 Data composition

To predict caRBPs, we adopted caRBPs and chromatin-contact peptides from mESCs (14). After filtering, we obtained a total of 385 caRBPs and 1630 non-caRBPs for training and testing (Figure 2A, Suppl. Table 1). Additionally, we compiled 1532 chromatin-contact peptides (defined as positive peptides) derived from caRBPs and 14,937 peptides (defined as negative peptides) from RBPs lacking chromatin-binding ability (Figure 2A-B, Suppl. Table 1). We extracted two key features from each protein or peptide sequence: the primary amino acid sequence and the predicted secondary structure information (Figure 2C, Suppl. Table 1), as primary sequences alone may not capture spatial and conformational aspects critical for chromatin binding.

**Figure 2.**
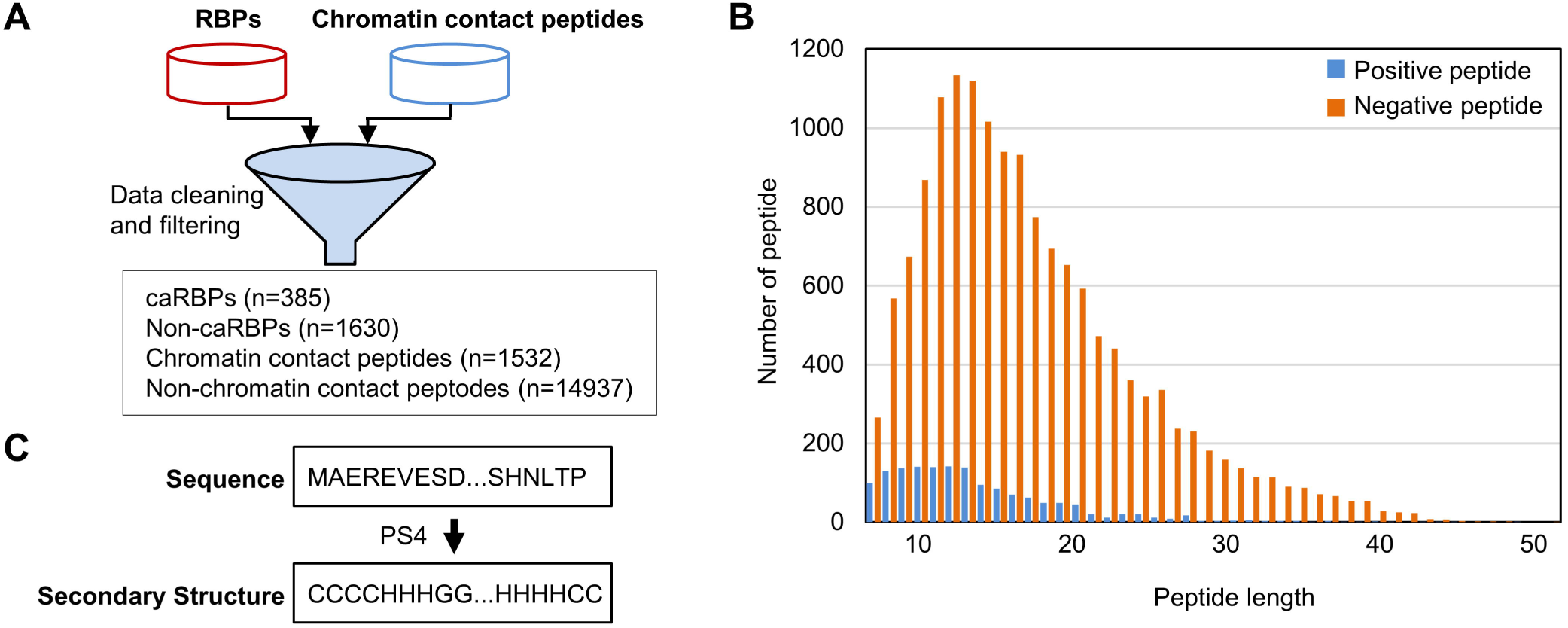
Data composition and processing. **(A)** Flowchart of the cleaning of raw data. **(B)** The distribution of peptide length was plotted. Distribution of protein length was not plotted because of the sparsity of length frequency (over half of proteins’ length frequency is 1). **(C)** Flowchart of the prediction of secondary structure.

### 3.2 CNN-BiLSTM model with chromatin-contact peptide can predict the caRBPs

Based on the model architecture described in the Methods, we constructed a CNN-BiLSTM model (Figure 1), comprising two CNN layers and two BiLSTM layers followed by dense and activation layers. We also constructed other learning models, including SVM, RF, and XGBoost, to evaluate the performance of each model on caRBP prediction (Figure 3A, Suppl. Table 3). The CNN-BiLSTM model achieved higher performance than other models, except for accuracy, where it was slightly lower than the Random Forest (RF) model (Figure 3B, Suppl. Table 3). Regarding feature sets, higher performance was observed for peptide sequence (pep seq) and the combination of peptide sequence and secondary structure (pep seq + SS). Upon further examination of the performance matrices on these two features, there is no significant difference between them. Taking all the performance metrics into consideration, the peptide sequence and combination of peptide sequence and peptide secondary structure were selected for further analysis with CNN-BiLSTM model.

**Figure 3.**
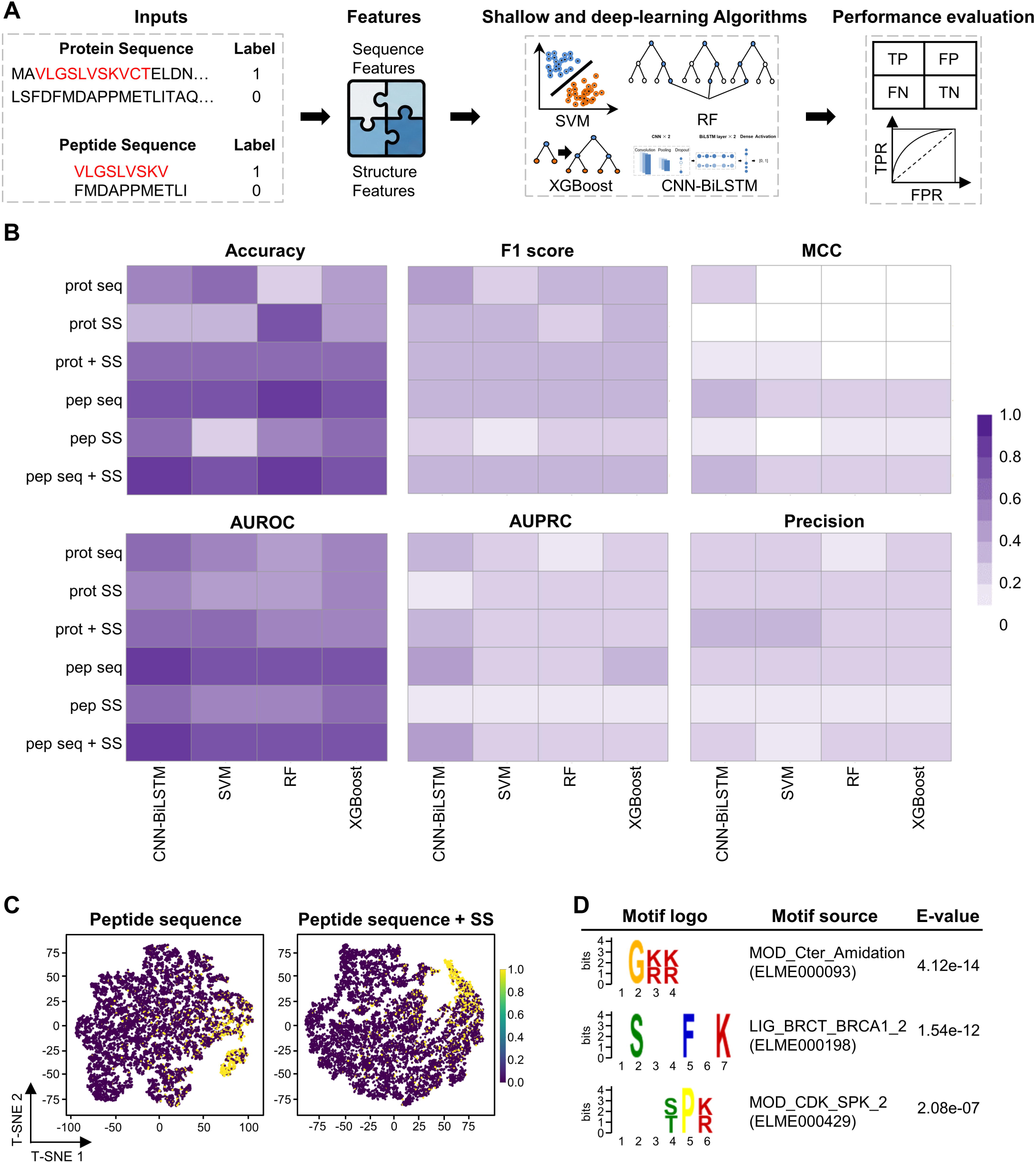
Evaluation of various model performance. **(A)** Flow diagram depicts the steps to develop the proposed caRBP prediction model. **(B)** Heatmap of the performance metrics of different learning algorithms with different features. **(C**) Feature visualization of CNN-BiLSTM model by t-SNE for dimension reduction. Feature representation of peptide sequence (left panel) and combination of peptide sequence and Secondary Structure (SS) (right panel). **(D)** XSTREME predicted protein motifs enriched in the chromatin-contact peptides.

Next, we extract the features from CNN-BiLSTM and feature visualization of peptide sequence or combination of peptide sequence and peptide secondary structure indicates that positive and negative peptides can be separated clearly, and there is no significant difference between them for both between-class scatter matrix and within-class scatter matrix (Figure 3C). These findings indicate that inclusion of peptide secondary structure information did not significantly enhance model performance.

To further investigate the intrinsic differences between positive and negative peptides, we performed motif enrichment analysis. We identified several motifs enriched in positive peptides, such as “.G[RK][RK]”, “.(S)..F.K”, and “…([ST])P[RK]” (Figure 3D). These motifs are associated with functional sites, including Peptide Amidation Sites, BRCA1 C-Terminus (BRCT) phosphopeptide ligands, and Cyclin-dependent kinases (CDKs) Phosphorylation Sites. Additionally, we compared the physicochemical properties of caRBPs and non-caRBPs using the cleverMachine algorithm (27). We found that nucleic acid binding ability could discriminate caRBPs (positive) from non-caRBPs (negative). In contrast, other properties such as disorder propensity, aggregation, burial propensity, and hydrophobicity could not (Supplementary Figure 1). These results may support that chromatin-contact peptides reflect the nucleic acid binding ability of caRBPs.

Collectively, these findings indicate that raw peptide sequences alone can accurately classify caRBPs and non-caRBPs, with nucleic acid binding ability of caRBPs being a key distinguishing feature.

### 3.3 Existing DRBP prediction tools fails to predict caRBP

We next examined whether previously mentioned DRBP predictors (iDRBP_MMC, DeepMC-iNABP, and DeepDRBP-2L) could accurately predict the caRBPs identified in mESCs. It is important to note that the source codes for these tools are not publicly available. Consequently, we evaluated their generalizability rather than conducting a direct comparison of model architectures under identical training conditions. Despite performing well on their original datasets, these tools displayed a very low recall rate on our caRBP dataset (iDRBP_MMC: 9/385, DeepMC-iNABP: 34/385, DeepDRBP-2L: 75/385).

### 3.4 caRBP-Pred can accurately predict the potential caRBPs in mice

Based on these results, we developed an in-house script named caRBP-Pred to predict potential caRBPs using positive and negative peptide sequences (Figure 4A, mentioned in Methods). To identify potential caRBPs in mice, we obtained a total of 2897 mouse RBPs from the RBP2GO database (23), excluding those already present in our training and testing datasets. This resulted in an independent dataset of 1559 RBPs (Figure 4B). Using caRBP-Pred, we predicted 52 out of these 1559 RBPs as potential caRBPs in mice (Figure 4B, Supplementary Table 4). Interestingly, known DRBP predictors only predict a limited number of them as caRBPs (iDRBP_MMC: 2/52, DeepMC-iNABP: 7/52, DeepDRBP-2L: 8/52, Suppl. Table 4). Although no direct experimental evidence confirmed the chromatin binding capabilities of these 52 caRBPs in mice, we examined their human homologs. Consulting the human protein-DNA interaction database hPDI (28), we identified 11 RBPs with confirmed chromatin binding capabilities (Figure 4C). Notably, caRBP-Pred accurately predicted 10 out of these 11 RBPs as potential caRBPs, except the ZCCHC8 (Figure 4D, Suppl. Table 4), achieving a high accuracy rate of 91%. Again, only two of these 11 caRBPs are correctly predicted by the known DRBP predictors (Suppl. Table 4). This underscores the reliability of our program in predicting caRBPs and highlights its potential utility in future studies.

**Figure 4.**
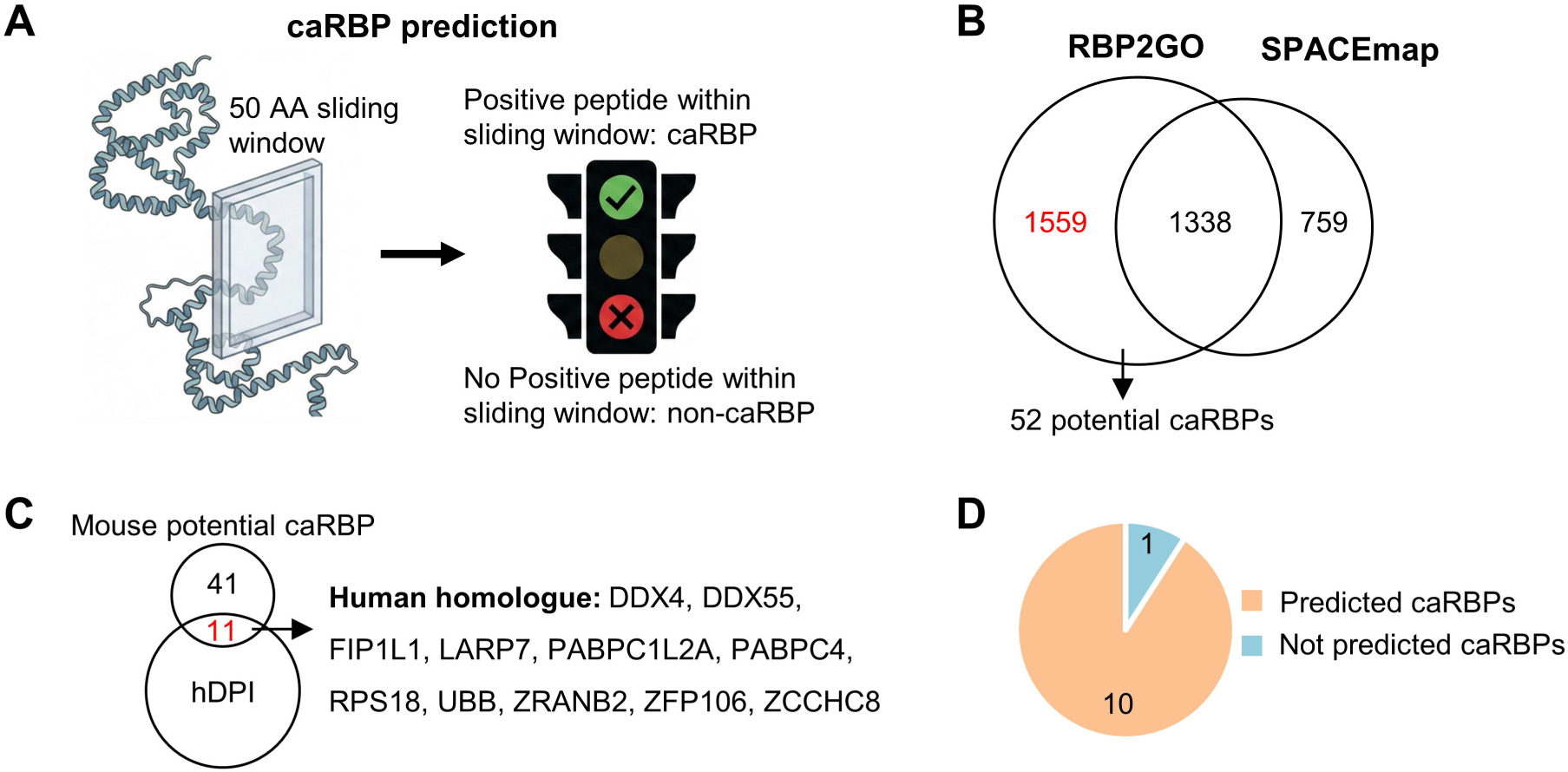
Prediction of caRBPs by caRBP-Pred. **(A)** Flowchart of caRBP-Pred. **(B)** Venn diagram shows the overlap between mouse RBPs from RBP2GO database and training and testing datasets from SPACEmap. A total of 1559 mouse RBPs were used for caRBP prediction, and 52 out of them were predicted as potential caRBPs. **(C)** Eleven RBPs of human homolog had experimental evidence to bind to chromatin. **(D)** Pie chart showed the number of predicted caRBPs of human homolog.

In conclusion, we constructed a CNN-BiLSTM model and developed caRBP-Pred to predict caRBPs based on chromatin-contact peptides. A total of 52 potential caRBPs were predicted in mice by using caRBP-Pred, and our caRBP-Pred showed better performance than the existing DRBP predictors. These predicted caRBPs can significantly help wet-lab researchers save time and funding, enabling them to directly explore the transcriptional or co-transcriptional role of caRBPs in their downstream experiments.

## 4 Discussion

Despite the widespread adoption of numerous computational methods for predicting RBPs, no tool has been specifically developed to predict this particular subtype of RBPs. In this study, we introduced a CNN-BiLSTM model for predicting caRBPs. This represents the first computational predictor capable of identifying mouse caRBPs.

Our CNN-BiLSTM model effectively captures local information and extracts intrinsic characteristics of caRBPs using only chromatin-contact peptide sequences, as evidenced by its high accuracy (AUC=0.81). Surprisingly, models trained on full-length protein sequences performed poorly. This suggests that the chromatin binding signal is enriched in specific domains, and using full-length sequences may dilute these critical features with irrelevant information. This finding aligns with the poor performance of existing DRBP predictors on our dataset, as they rely on full-length sequence inputs. This tool is particularly valuable for assisting researchers in predicting and uncovering caRBPs from a vast array of RBPs with unknown chromatin-binding capabilities. It holds significant potential for elucidating the regulatory roles of caRBPs in gene expression.

It is important to note that, despite the widespread use of CNN-LSTM and CNN-BiLSTM architectures in RBP prediction, these methods have several drawbacks, such as long training times and the requirement for high processing power (29). Other algorithms, such as transfer learning, stacking, and multilayer perceptron, have also been employed for RBP prediction (30,31). In fact, we initially attempted to use a transformer-based model for training. However, due to the limited sample size, this architecture was not well-suited for the current task. Given that the primary advantage of transformers is the ability to process entire input sequences simultaneously, such models could be highly effective for predicting caRBPs once larger peptide sequence datasets become available.

In summary, our CNN-BiLSTM model successfully predicted caRBPs using only chromatin-contact peptide sequences, and caRBP-Pred provided a valuable list of potential caRBPs in mice.

## Data availability

The datasets, source code, and user notes are available on GitHub (https://github.com/callmeracle/caRBP-Pred).

## CRediT authorship contribution statement

**Qiang Sun:** Writing – original draft, Conceptualization, Methodology, Formal analysis, Data curation. **Feng Yang:** Methodology. **Hao Sun:** Writing – original draft, Writing – review & editing, Supervision, Project administration, Funding acquisition. **Xiaona Chen:** Writing – original draft, Writing – review & editing, Supervision, Project administration. **Huating Wang:** Writing – review & editing, Writing – original draft, Conceptualization, Supervision, Resources, Project administration, Funding acquisition.

