## Supplemental Information for "caRBP-Pred: Deep Learning-based Prediction of Chromatin-Associated RNA-Binding Proteins Using Short Peptide Sequences"

**Inventory of Supplementary Information**

**1. Supplementary Figures**

Figure S1. Physico-chemical properties of caRBPs (positive) and non-caRBPs (negative) predicted by cleverMachine.

**2. Supplementary Tables**

Supplementary Table S1. Data utilized in this study.

Supplementary Table S2. caRBP prediction results from existing computational tools.

Supplementary Table S3. Evaluation of performance of various models and input features.

Supplementary Table S4. Predicted potential mouse caRBPs and the performance of existing DRBP predictors on these candidates.

**3. Supplementary Figures Legends**

**Supplementary Figure S1. Physico-chemical properties of caRBPs (positive) and non-caRBPs (negative) predicted by cleverMachine.** Bar plots were generated by the cleverMachine web server based on the input of caRBPs and non-caRBPs sequences.
