## Supplementary figures and images for "caRBP-Pred: Deep Learning-based Prediction of Chromatin-Associated RNA-Binding Proteins Using Short Peptide Sequences"

### Supplemental Figure

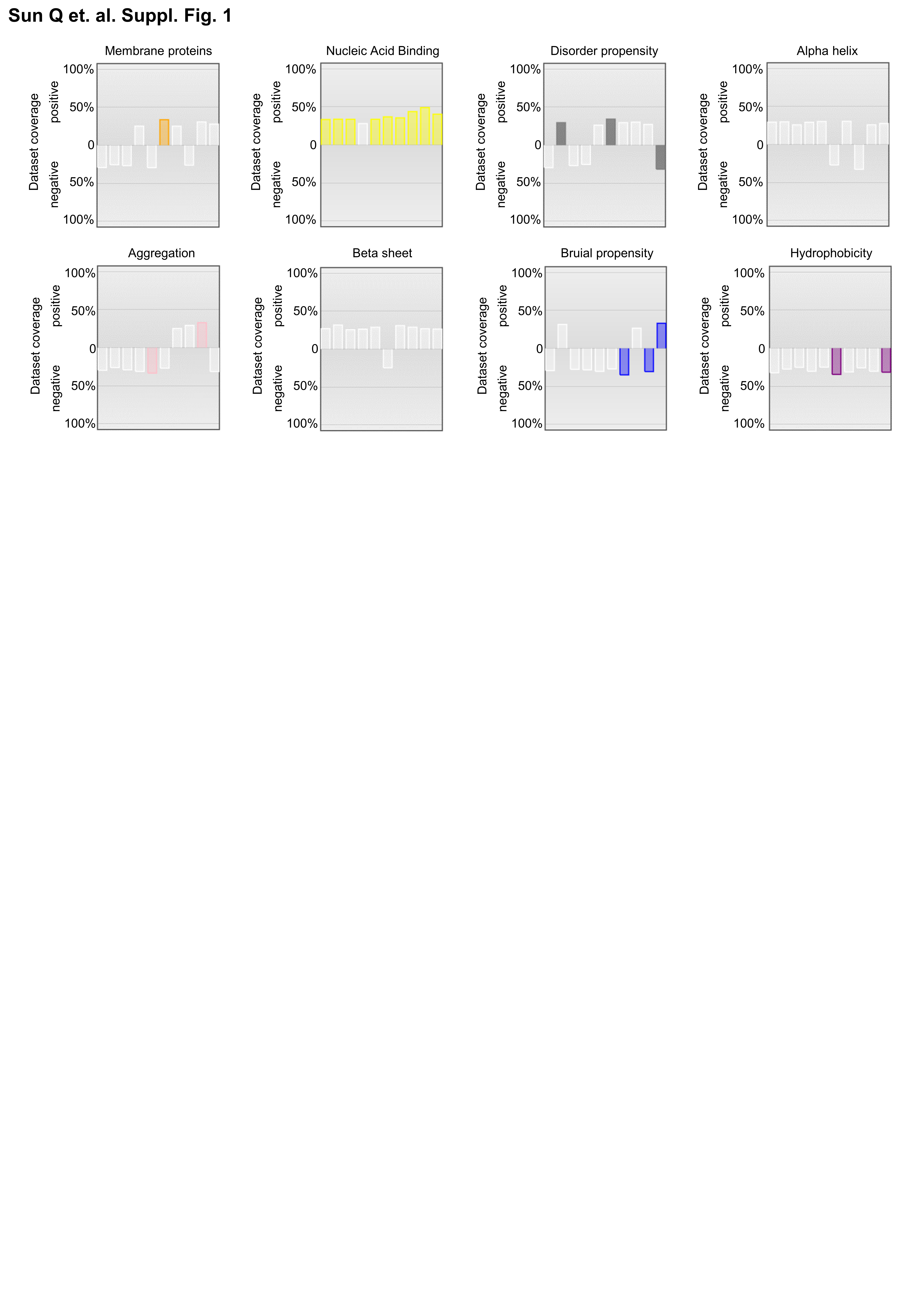
